# MNRR1 activation by nitazoxanide abrogates lipopolysaccharide-induced preterm birth in mice

**DOI:** 10.1101/2023.01.30.525762

**Authors:** Neeraja Purandare, Nardhy Gomez-Lopez, Marcia Arenas-Hernandez, Jose Galaz, Roberto Romero, Yue Xi, Andrew Fribley, Lawrence I. Grossman, Siddhesh Aras

## Abstract

Intra-amniotic inflammation leading to preterm birth is one of the leading causes of neonatal morbidity and mortality. We recently found that the mitochondrial levels of MNRR1 (Mitochondrial Nuclear Retrograde, Regulator 1; also called CHCHD2, AAG10, or PARK22), an important bi-organellar regulator of cellular function, are reduced in the context of inflammation and that both genetic and pharmacological increase in MNRR1 levels can counter the inflammatory profile. We show here that nitazoxanide, a clinically-approved drug, is an activator of MNRR1 and prevents preterm birth in a well-characterized murine model caused by intra-amniotic lipopolysaccharide (LPS) injection.

**Highlights:**

- Nitazoxanide exerts anti-inflammatory functions via activation of MNRR1.
- Oral administration of nitazoxanide prevents preterm birth in mouse model of intra-intraamniotic LPS-induced inflammation.

## Introduction

Intra-amniotic inflammation leading to preterm birth is one of the leading causes of neonatal morbidity and mortality [1-3]. Intra-amniotic inflammation results from ascending microbial invasion from the lower genital tract to the amniotic cavity, termed intra-amniotic infection, or from endogenous danger signals (i.e., alarmins) released upon cellular damage or stress, known as sterile intra-amniotic inflammation [3-5]. Although immune cells are known to play a key role in the response to inflammation at the fetal-maternal interface [6,7], the specific mechanisms underlying the response of cells in the placental tissues to inflammatory insults are largely unknown [8]. We recently identified a non-canonical, TLR4-independent signaling pathway involving mitochondria in placental cells that contributes to immune pathology [9]. Using both *in vitro* and *in vivo* models, we found that the mitochondrial levels of MNRR1 (Mitochondrial Nuclear Retrograde, Regulator 1; also called CHCHD2, AAG10, or PARK22), an important bi-organellar regulator of cellular function [10-14], are reduced in the context of inflammation. However, whether MNRR1 is implicated in the pathophysiology of intra-amniotic inflammation-induced preterm labor and birth has not been well explored.

MNRR1 functions in two cellular compartments, the mitochondria and the nucleus. Mitochondrial MNRR1 binds to cytochrome *c* oxidase [11] or Bcl-xL [15] to regulate the mitochondrial roles of energy generation and apoptosis, respectively. Nuclear MNRR1 can function as a transcriptional regulator to modulate the activation of stress-responsive genes including MNRR1 itself [11,16]. This novel LPS-induced inflammatory signaling pathway was characterized *in vitro* using a human trophoblast cell line (HTR8/SVneo) [9]. The pro-inflammatory pathway is initiated by activation of NOX2, generating radical oxygen species (ROS) to activate ATM kinase that then phosphorylates YME1L1, a mitochondrial inter-membrane space protease, which subsequently reduces mitochondrial MNRR1 levels. Furthermore, overexpression of MNRR1 was shown to dampen this inflammatory signaling pathway and restore mitochondrial function.

Given the observed beneficial effects of restoring MNRR1 expression *in vitro*, we recently performed a screen of FDA-approved drugs to identify pharmacological activators of MNRR1 and were able to identify nitazoxanide as an inducer of MNRR1 transcription. Moreover, we showed that nitazoxanide could rescue the trophoblast phenotype induced by LPS treatment [9]. Therefore, in the current study we examined whether nitazoxanide could represent a viable strategy of preventing preterm birth induced by a microbial product.

## Results

### The anti-inflammatory effects of nitazoxanide are mediated via transcriptional activation of MNRR1

Nitazoxanide was identified as an MNRR1 activator from a list of 2,400 FDA-approved compounds in a high-throughput screen (Figure 1A). By using orthogonal assays with HTR8 cells, we found that the half-maximal effective concentration (EC50) of nitazoxanide is 0.115 µM (Figure 1B). To evaluate whether the anti-inflammatory effects of this compound are mediated via activation of MNRR1, we tested the MNRR1 dependence of the ability of nitazoxanide to mitigate the LPS-induced inflammatory response in HTR8 cells. We found that activation of *TNF* transcription, a previously described effect of LPS exposure in HTR8 cells [9], was reduced to control levels in an MNRR1-dependent fashion (Figure 1C). There was also a basal increase in *TNF* transcription that was restored to control levels upon reintroduction of MNRR1. The cause of this increase is yet to be clarified and will be explored in future studies.

**Figure 1.**
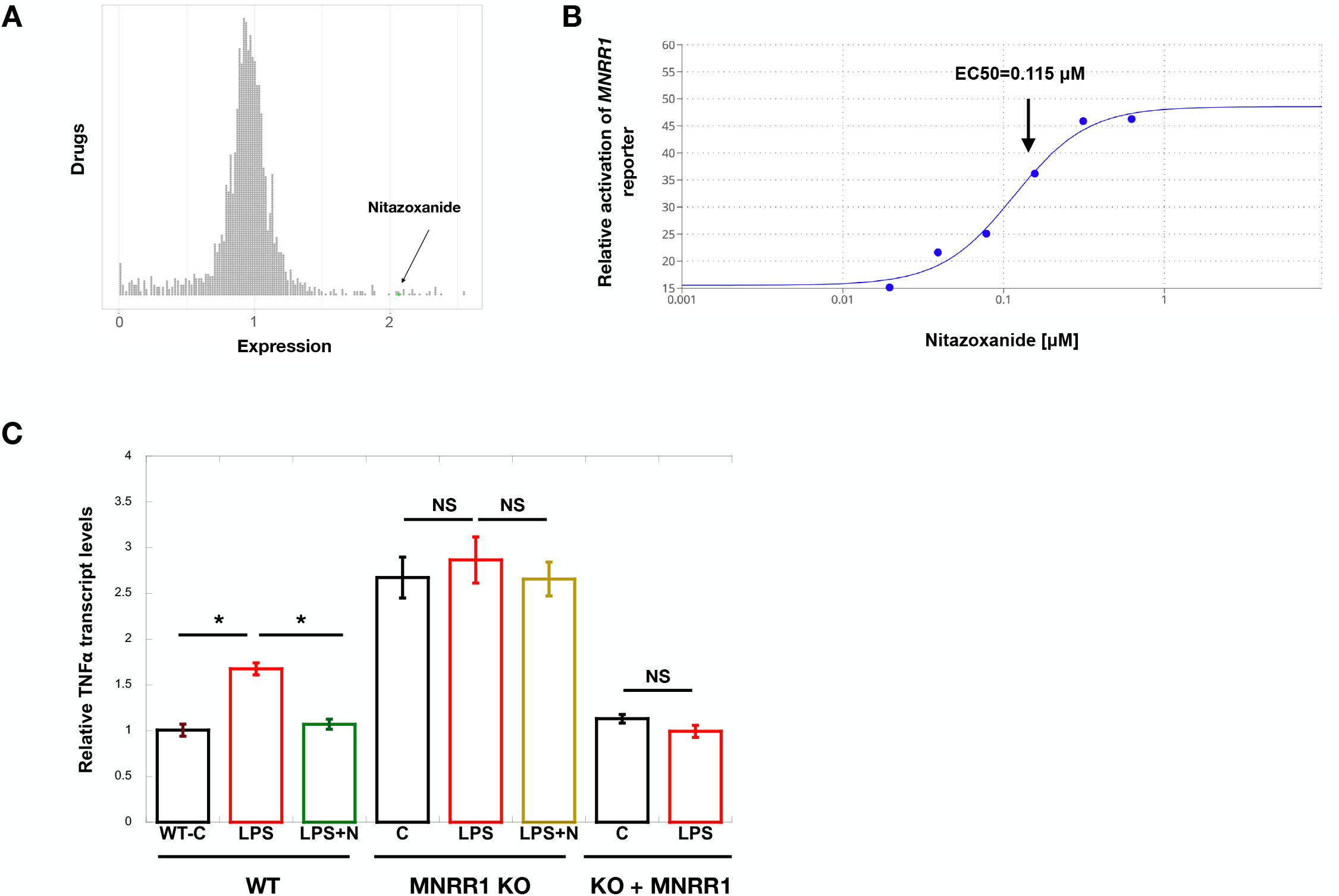
Nitazoxanide’s anti-inflammatory effects are via transcriptional activation of MNRR1. **A:** Results of a screen of 2,400 FDA-approved drugs identified to transcriptionally activate (>1), inhibit (<1), or not affect (=1) MNRR1. Each circle represents one drug and the MNRR1 activator (Nitazoxanide (N)) has been highlighted in green. **B:** Equal numbers of HTR cells overexpressing the MNRR1-luciferase reporter were plated on a 96 well plate and treated with increasing amounts of nitazoxanide or vehicle (DMSO) for 24 h. EC50 was calculated using the activation of MNRR1 reporter relative to vehicle treated cells using the Quest Graph(tm) EC50 Calculator. **C:** *TNF* transcript levels were measured in human placental cells. Actin was used as a housekeeping gene to normalize expression (n=4, for all figures, significance is indicated with * and represents p-value<0.05, ns is non-significant).

### Activation of MNRR1 using nitazoxanide prevents preterm birth in vivo

We next tested the effects of nitazoxanide in a well-characterized murine model of intra-amniotic LPS injection that results in inflammation and preterm birth [17-20]. Dams received oral administration of nitazoxanide and ultrasound-guided intra-amniotic injection of LPS as shown in Figure 2A. Nitazoxanide extended the gestational age, preventing LPS-induced preterm birth [delivery <18.5 days *post coitum* (dpc)] in all pregnant dams (Figure 1B). Furthermore, neonatal mortality at birth was also significantly reduced in dams treated with nitazoxanide compared to controls (Figure 2C). Finally, we showed that the LPS-induced reduction of MNRR1 was restored at the transcript level in the decidua (Figure 2D) and at the protein level in the placenta (Figure 2E) by nitazoxanide treatment.

**Figure 2.**
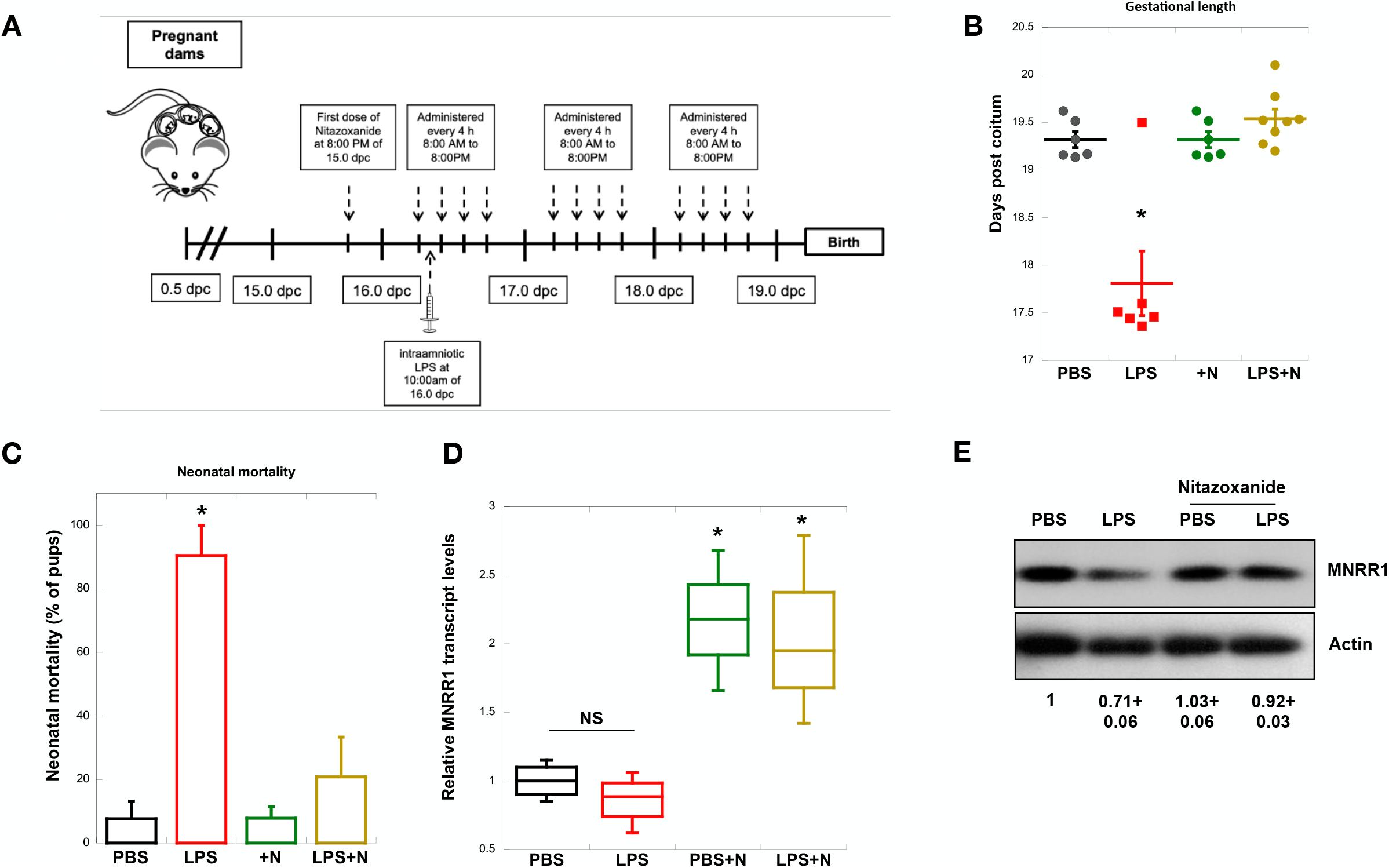
Nitazoxanide prevents preterm delivery and improves neonatal mortality *in vivo*. **A:** Schematic illustration depicting the administration of nitazoxanide (Alinia) and LPS in the murine model of preterm birth. **B:** Gestational length of mice was obtained using video-monitoring until delivery (n=6-8). **C:** Neonatal mortality was determined by calculating the number of dead pups over total litter size (n=6-8). **D:** MNRR1 transcript levels were measured in mouse decidual tissue (17.5 dpc). Actin was used as a housekeeping gene to normalize expression (n=4 for all except PBS+N, where n=3). **E:** Equal amount of whole placental lysates were separated on an SDS-PAGE gel and probed for MNRR1 levels. Actin was used as a loading control (n=2).

## Discussion

We show here that nitazoxanide, a clinically-approved drug sold commercially as Alinia, can prevent preterm birth in an LPS-based mouse model of intra-amniotic inflammation by stimulating the transcription of MNRR1. We previously showed that LPS treatment produces an inflammatory phenotype in a trophoblast cell line that includes reduced mitochondrial respiration and increased production of ROS and inflammatory cytokines such as TNF [9]. This phenotype included reduced expression of the mitochondrial regulator MNRR1, and stimulation of this factor either genetically or pharmacologically with nitazoxanide was able to reverse the inflammatory profile. Consistently, we show here that the anti-inflammatory effects of nitazoxanide in pregnant mice are also mediated through transcriptional activation of MNRR1.

Nitazoxanide is currently used as a treatment for several infectious organisms including protozoa, helminths, bacteria, and viruses [21]. The molecular target of nitazoxanide is known in protozoa [22], but not in humans. Inflammation-induced preterm birth leads to significant adverse effects on maternal and fetal health [23]. Nitazoxanide, a clinically approved drug that is safe to use during pregnancy [24] and can be administered orally, may be a useful addition to evaluate for high-risk pregnancies.

The mechanisms by which nitazoxanide exerts its anti-inflammatory effects in mice will require further study. MNRR1 is known to function in both mitochondria and the nucleus [10-14], and in a recent cellular model its nuclear function alone as a transcriptional activator was sufficient to dampen LPS-induced inflammation [9]. However, in animals a more detailed understanding of its action in activating multiple homeostatic mechanisms [16] may well uncover additional genes of interest as well as its potential utility for treating other inflammatory conditions.

## Materials and methods

### Methods details

#### Cell culture and reagents

##### Cell lines

All cell media were supplemented with 5% fetal bovine serum (FBS) (Sigma Aldrich, St. Louis, MO, USA) plus Penicillin-Streptomycin (HyClone, Logan, UT, USA). HTR8/SVneo (HTR) cells were cultured in Roswell Park Memorial Institute Medium (RPMI) (HyClone, Logan, UT, USA).

##### Chemicals

Nitazoxanide was obtained from Selleckchem (Houston, TX, USA) and solubilized in DMSO (used as vehicle control in all nitazoxanide experiments). Ultrapure LPS for cell culture experiments (lipopolysaccharide from *Escherichia coli* 0111:B4) was purchased from Invivogen (San Diego, CA, USA).

##### Plasmids

The *MNRR1* promoter firefly luciferase reporter plasmid and pRL-SV40 (*Renilla*) luciferase plasmids (for normalization of firefly luciferase) have been described previously [11]. All plasmids were purified using the EndoFree plasmid purification kit from Qiagen (Germantown, MD, USA).

#### Transfections and luciferase reporter assays

HTR cells were transfected with the indicated plasmids using TransFast transfection reagent (Promega,Madison, WI, USA) according to the manufacturer’s protocol as previously described [9]. Luciferase assays were performed with the dual-luciferase reporter assay kit (Promega, Madison, WI, USA). Transfection efficiency for the MNRR1 reporter was normalized with the co-transfected pRL-SV40 *Renilla* luciferase expression plasmid [10,12,13]. EC50 was calculated using Quest Graph(tm) EC50 Calculator (*AAT Bioquest, Inc*., https://www.aatbio.com/tools/ec50-calculator).

#### Real-time polymerase chain reaction

Total cellular RNA was extracted from mouse placental tissues or HTR8 cells using a RNeasy Plus Mini Kit (Qiagen, Germantown, MD, USA) according to the manufacturer’s instructions. Complementary DNA (cDNA) was generated by reverse transcriptase polymerase chain reaction (PCR) using the ProtoScript® II First Strand cDNA Synthesis Kit (New England Biolabs, Ipswich, MA, USA). Transcript levels were measured by real time PCR using SYBR green on an ABI 7500 system. Real-time analysis was performed by the ΔΔ^Ct^ method. The primer sequences used were as follows: *MNRR1* mouse (forward) - 5’-ATGGCCCAGATGGCTACC-3’, *MNRR1* mouse (reverse) - 3’-CTGGTTCTGAGCACACTCCA-5’; Actin mouse (forward) -5’-TCCTCCCTGGAGAAGAGCTA-3’, Actin mouse (reverse) - 3’-ACGGATGTCAACGTCACACT-5’, *TNF* human (forward) - 5’-TGTAGCAAACCCTCAAGCTG-3’ *TNF* human (reverse) 3’-GAGGTTGACCTTGGTCTGGT-5’, Actin human (forward) -5’-CATTAAGGAGAAGCTGTGCT-3’, Actin human (reverse) - 3’-GTTGAAGGTAGTTTCGTGGA-5’

#### Immunoblotting

Immunoblotting was performed as previously described [10,11]. Tissue lysates were prepared using 50 mM Tris-HCl (pH 8.0) with 150 mM NaCl, 1% NP-40, 0.5% sodium deoxycholate, and 0.1% SDS. All lysis buffers included a protease and phosphatase inhibitor cocktail (Sigma, PPC1010).

#### Mice

C57BL/6 mice were purchased from The Jackson Laboratory (Bar Harbor, ME, USA), and bred in the animal care facility at the C.S. Mott Center for Human Growth and Development at Wayne State University (Detroit, MI, USA). Mice were housed under a circadian cycle (12 h light:12 h dark). Eight-to twelve-week-old females were mated with males of proven fertility. Female mice were examined daily between 8:00 a.m. and 9:00 a.m. for the presence of a vaginal plug, which indicated 0.5 days *post coitum* (dpc). Female mice with a vaginal plug were removed from the mating cages and housed separately. A weight gain of ≥2 grams confirmed pregnancy at 12.5 dpc. All animal experiments were approved by the Institutional Animal Care and Use Committee at Wayne State University (Protocol No. 21-04-3506).

#### Animal model of intraamniotic LPS-induced preterm labor and birth

Dams were anesthetized on 16.5 dpc by inhalation of 2–3% (Fluriso(tm) (Isoflurane, USP) Vetone Boise, ID, USA) and 1–2 L/min of oxygen in an induction chamber. Anesthesia was maintained with a mixture of 1.5–2% isoflurane and 1.5–2 L/min of oxygen. Dams were positioned on a heating pad and stabilized with adhesive tape. Fur removal from the abdomen and thorax was performed by applying Nair cream (Church & Dwight Co., Inc., Ewing, NJ, USA) to those areas. Body temperature was detected with a rectal probe (VisualSonics Inc., Toronto, Ontario, Canada) and maintained in the range of 37±1°C. Respiratory and heart rates were monitored by electrodes embedded in the heating pad. An ultrasound probe was fixed and mobilized with a mechanical holder, and the transducer was slowly moved towards the abdomen. Ultrasound-guided intra-amniotic injection of lipopolysaccharide of *Escherichia coli* O111:B4 (LPS; Sigma-Aldrich, St. Louis, MO, USA) at a concentration of 100 ng dissolved in 25 µL of sterile 1X PBS was performed into each amniotic sac using a 30 G needle (BD PrecisionGlide Needle), and controls were injected with 25 µL of sterile 1X PBS alone. The syringe was stabilized by a mechanical holder (VisualSonics Inc.). Following ultrasound, dams were placed under a heat lamp for recovery (dam regaining normal activity, such as walking and responding), which occurs 10 - 15 min after removal from anesthesia. Following injection, dams were monitored using a video camera with infrared light (Sony) until delivery. Video monitoring allowed for determination of gestational length, which was calculated from the presence of the vaginal plug (0.5 dpc) until the observation of the first pup in the cage bedding. Preterm birth was defined as delivery before 18.5 dpc, and its rate was represented by the percentage of dams delivering preterm among the total number of mice injected. The rates of neonatal mortality at birth were calculated as the number of pups found dead among the total litter size.

#### Nitazoxanide treatment

Dams were treated with 200 mg/kg of nitazoxanide (Alinia, ROMARK, Tampa, FL, USA) via oral gavage at 8 pm on 15.5 dpc, again at 8 am, 12 pm, 4 pm, and 8 pm on 16.5 dpc, and then the tissues were collected the morning of 17.5 dpc for molecular experiments. These dams were intra-amniotically injected with either LPS or PBS control at 10 am on 16.5 dpc.

